# Unveiling Convergent and Divergent Intrinsic Brain Network Alternations in Depressed Adolescents Engaged Non-Suicidal Self-Injurious Behaviors with and without Suicidal Attempts

**DOI:** 10.1101/2023.03.09.531863

**Authors:** Linling Li, Zhen Liang, Guohua Li, Hong Xu, Xing Yang, Xia Liu, Xin Zhang, Jianhong Wang, Zhiguo Zhang, Yongjie Zhou

## Abstract

Nonsuicidal self-injury (NSSI) and suicidal attempt (SA) frequently occur during depressed adolescents, but the underlying neurobiological mechanisms are poorly understood. The maturation of network connectivity is a central feature of brain development during adolescence. However, few researchers have used resting-state functional magnetic resonance imaging (fMRI) to investigate the similarities and differences in the intrinsic brain networks between NSSI with NSSI+SA in depressed adolescent populations. In this study, we collected self-reported questionnaires and resting-state (fMRI data from a sample of depressed adolescents (N = 114), categorized into three groups: clinical control (non-self-harm), NSSI-only, and NSSI+SA based on their self-harm history. The alternations of FC patterns were identified through support vector machine-based classification, as machine learning approaches can help to overcome the multiple-comparison problems of their multivariate nature. Referring to the convergent alternations between adolescent NSSI with or without a history of SA, the alternations of intra-/inter-network FCs among the Control, SalVAtten, Limbic, and Default networks. Besides, divergent alternations were also observed for depressed adolescents with a history of SA, involving the Limbic, DorsAttn, Visual, and Subcortical networks. Besides, the severity of depressive symptoms only showed a significant correlation with altered FCs in Limbic-DorAttn and Limbic-Visual networks, strengthening the fact that these observed alternations of FC could not all be explained by increased depression severity. The emphasis of this study on the network basis could extend the existing evidence at a circuit level and the observed convergent alternations may explain an underlying neurobiological mechanism pertaining to the continuum of self-destructiveness in adolescents. In addition, our discovery of divergent alternations could help to identify biomarkers that will aid in differentiating those at risk for SA among those engaging in NSSI and contribute to the development of neurobiologically informed interventions.

## INTRODUCTION

Adolescence, a period characterized by significant physiological, psychological, and social changes [1], poses a heightened risk for mental disorders and maladaptive behaviors, such as depression and self-injury behaviors. Nonsuicidal self-injury (NSSI) refers to intentional, self-inflicted damage to the surface of the body without intent to kill oneself, which is not socially sanctioned [2]. On the other hand, suicidal attempts (SA) involve engaging in potentially self-injurious behavior with some intent to die [3]. Both NSSI and SA share common heritable factors [4] and high levels of comorbidity [5] and often co-occur during adolescence, especially in clinical samples [6, 7]. Adolescents may initially experience depression and subsequently develop NSSI as a maladaptive coping mechanism to alleviate emotional pain [8]. The prevalence rates of NSSI among the general population are estimated to be about 17% in community samples [9] and more than 40% in clinical samples [10]. Although not all self-injurers will attempt suicide, a high frequency of NSSI incidents is associated with an increased risk of SA [11]. Adolescents with repeated NSSI who additionally reported a history of SA often exhibit greater severity of psychopathology compared to those with NSSI alone [12]. Given that SA is associated with more challenging treatment courses and an elevated risk of mortality [13], it is important to understand why certain adolescents only engage in NSSI whereas others engage in SA. Investigating the neurobiological underpinings of these behaviors during adolescence is particular important, as it can inform early intervention and prevention strategies that promote healthy neurodevelopmental trajectories [14].

To date, a limited number of neuroimaging studies have investigated NSSI and SA in adolescents, as discussed in comprehensive reviews [15–17]. These studies have revealed a complex and diverse array of neural mechanisms underlying these behaviors. Specifically, research on youth SA has consistently identified alternations in the prefrontal and limbic systems, suggesting that a comprised regulatory system may potentiate suicidal thinking, while perturbations in both top-down and bottom-up connections may lead to SA [15, 16]. In the realm of youth NSSI research, the primary focus has been on disruptions in the amygdala and anterior cingulate cortex activations, as well as functional connectivity (FC) patterns, which contribute to hyperarousal and reduced impulse control-center features of NSSI [16]. Among these studies, the majority were conducted with task-based fMRI and regions-of-interest (ROI) based FC analysis. While these task-specific connectivity findings are important in beginning to elucidate the brain basis, these studies utilized disparate emotion processing tasks and obtained discrepant findings, signaling the need for understanding intrinsic (i.e., task-independent) neural networks associated with NSSI and SA. On the other hand, further exploration of task-independent networks could obtain a more holistic view of the neurobiology of NSSI and SA. Such knowledge will not only fill existing gaps but also pave the way for further research endeavors and inform the development of targeted interventions.

Resting-state fMRI analysis has emerged as a valuable tool for investigating intrinsic large-scale functional networks [18], and it offers significant advancements in our understanding of whole-brain network-based alternations in clinical conditions. Task-independent patterns of intrinsic functional signals derived from resting-state fMRI studies are relatively stable and have strong test re-test reliability [19]. However, compared with structural and task-based fMRI studies, the resting-state fMRI literature on adolescent NSSI and SA is smaller, and interpretations are challenging, given mixed methods and results [16]. Many of the existing studies in this area are hypothesis-driven and primarily focused on selected ROIs, such as the amygdala [20] and posterior cingulate cortex [21]. Although one recent study applied a data-driven approach (e.g. independent component analysis) to explore brain networks like the default mode network [22–26], a comprehensive whole-brain analysis is necessary to expand the scope of investigated brain networks. Such a whole-brain analysis would enable a more complete understanding of neurobiological underpinnings associated with NSSI and SA in adolescents.

Investigating the clinical phenomena of NSSI and suicidal behavior poses challenges due to their independent and co-occurring nature in adolescence [27]. According to the proposed Gateway Theory, suicidal behaviors stem from escalating NSSI behaviors [28]. A previous study examined the neural correlates of NSSI and suicidal ideation (SI) based on an overlapping sample of depressed adolescents, distinguishing between those with or without a history of NSSI (SI) [22]. Their findings revealed that both SI and NSSI were related to brain networks associated with difficulties in self-referential processing and future planning difficulties, while NSSI specifically was related to brain networks associated with disruptions in interoceptive. It has been reported that a third of adolescents with suicidal thoughts eventually progress to suicide attempts [29], with NSSI behavior playing a specific role in the transition from thoughts to actions [30]. Consequently, there is a need for studies that explore neurobiological correlates capable of differentiating between adolescents engaged in NSSI who attempt suicide and those who do not, thereby enhancing our current knowledge and informing clinical practice. There may exist a complex neurobiological interaction and overlap in resting-state FC patterns between NSSI and SA. However, so far, few researchers have used resting-state fMRI to investigate the similarities and differences in the intrinsic brain networks between NSSI with NSSI+SA in depressed adolescent populations.

The present study examined NSSI and/or SA within the context of depression. We specifically examined the presence of the previous history of SA among adolescents presenting with NSSI, aiming to uncover both convergent and divergent patterns of self-harm behaviors (NSSI only, NSS+SA) when compared with clinical controls. Findings based on a comparison between diagnostic controls and individuals with the same mental disorder plus NSSI or SA are more commonly reported in the literature and are more likely to reflect specific effects of this behavior in that disorder. Whole brain resting-state FC alternations were extracted using data-driven support vector classification analysis, as machine learning approaches can help to overcome the multiple-comparison problems of their multivariate nature [31]. Based on the literature outlined above, we hypothesize that both convergent and divergent patterns of resting-state FC alternations exist for different patterns of self-harm behaviors (NSSI only, NSS+SA) in depressed adolescents. A thorough exploration of the neural correlates underlying NSSI and SA may lead to a more comprehensive understanding of the nature of the link between these two types of self-destructive behavior in adolescents.

## MATERIALS AND METHODS

### Participants

Depressed adolescents were recruited from the Department of Depression at Shenzhen Kangning Hospital. Inclusion criteria were as follows: (1) age 12 to 18 years, (2) diagnosed with current depression by senior psychiatrists according to the Diagnostic Criteria and Statistical Manual of Mental Disorder Fifth Edition (DSM-5). The exclusion criterion consisted of a history of any serious neurological or intellectual disability, contraindications to MRI scanning, and meeting for a lifetime or current criteria for Mania, Psychosis, or any substance use disorders. In this study, 114 adolescents were recruited and completed the MRI data collection. The protocol for this study was approved by the Research Ethics Committee of Shenzhen Kangning Hospital (No: 2020-K021-04-1). Informed and written consent was obtained from both the participants themselves and their parents or caregivers.

The Chinese version of the Functional Assessment of Self-Mutilation (FASM) [32] was used to assess methods, frequency, and functions of NSSI behaviors in the past year (for details of the assessment content and score calculation, see supplementary material 1.1). Patients were categorized into three groups: (1) depressed adolescents without NSSI and SA serving as the diagnostic control group, (2) depressed adolescents with NSSI but without a history of SA, referred as the NSSI group, (3) depressed adolescents with a history of both NSSI and SA, referred as the NSSI+SA group. Considering the well-established link between emotion regulation deficits and self-injury behavior, all the enrolled adolescents completed the Chinese version of ERQ-CA [33] to access individual differences in the habitual use of two emotion regulation strategies.

### Data acquisition and preprocessing

MRI was collected and the data preprocessing was performed using Data Processing Assistant for Resting-State fMRI Advanced Edition V2.2 (DPARSFA, http://rfmri.org/DPARSF) toolkit with Statistical Parametric Mapping (SPM8, https://www.fil.ion.ucl.ac.uk/spm/) (for details of fMRI acquisition and processing, see supplementary material 1.2).

### Functional network construction

Whole brain resting-state FC matrices were calculated based on Pearson’s correlation in eight distinct brain networks given by the Schaefer cortical atlas with 200 parcels [34] and Melbourne Subcortical Atlas with 32 parcels [35] (supplementary material 1.3 and Figure S1). The feature vector of resting-state FC was constructed by vectorizing the upper triangular part of the correlation matrix of each subject.

### Support vector machine training and classification

Given the relatively high dimensionality of whole-brain brain network features, the utilization of machine learning approaches becomes crucial in addressing the multiple-comparison problem inherent in such multivariate data [31]. In this study, the identification of differential resting-state FCs between groups was accomplished through the application of a support vector machine (SVM)-based classification method. SVM analysis at the group level offers a powerful tool for identifying informative voxels or ROIs that contribute to the classification of different groups, allowing for the effective integration of information across multiple spatial locations. In accordance with previous studies using machine-learning methods for brain imaging classification [36], the original multiclass problem was translated into a series of binary comparisons, which means the linear binary SVM classifier [37] was trained using the feature sets derived from resting-state FC strength to classify each pair of groups. To ensure robust and reliable classification results, a nested leave-one-out-cross-validation (LOOCV) strategy was implemented. This approach helps to mitigate overfitting and provides a more accurate estimate of the classifier’s performance. Detailed information regarding the implementation of the SVM-based classification and the nested LOOCV procedure can be found in supplementary material 1.4.

### Evaluating contributions to a predictive model

By employing linear kernel SVM classifiers, it becomes possible to extract weights that reflect the significance of each feature for classification purposes, thereby allowing for the identification of group-level differences. Consensus features were defined as the common features always selected to form the final feature set for each LOOCV iteration. The weights of the consensus features were the average value of the classification weight across all iterations of LOOCV (weights of selected features were normalized to [–1 1], as shown in Figure S2). Then we only focused on the consensus features that exhibited at least one standard deviation greater weights than others. This selection criterion ensured that only those features with the most substantial discriminative power were prioritized for future analysis.

To increase the interpretability of the network-level contributions, we examined the modular architecture of these consensus features at the brain location level. The 232 ROIs can be divided into eight modules (intrinsic brain networks: Visual, somatomotor [SomMot], dorsal attention [DorsAttn], salience/ventral attention [SalVAttn], Limbic, Control, Default, Subcortical). To assess the contributions of these modules to the classification of different groups, we calculated the mean classification weights and percentages of intra- and inter-modular discriminating features for each group. For each subject, the mean weights were calculated by averaging the weights of all the discriminating features (consensus features with greater weights) across all LOOCV loops within each module and between pairs of modules. Furthermore, we also calculated the percentage (density) of discriminating features occurring within and between different modules (number of discriminating connections divided by the number of all possible connections). To focus on the most informative modular patterns, we specifically considered the discriminating intra- and inter-modular discriminating features that exhibited at least one standard deviation greater percentage than the remaining features (distribution density).

The predictive contributions of the consensus intra- and inter-modular features with both greater classification weights and densities were also visualized at the nodal level. Specifically, the weight of each region was calculated by summing one-half of the feature weights associated with that region [38]. Following this, the anatomical locations of the brain regions that play a more important role in the classification can be identified.

### Statistical analysis

All statistical analyses were performed using SPSS (IBM, Armonk, NY). The Chi-square test and one-way analysis of variance (ANOVA) test were performed to analyze the group differences in the demographic data. P < 0.05 was considered statistically significant.

The statistical significance of selected classifiers was evaluated by the non-parametric permutation tests (5000 times). In each permutation, we randomly assigned the group labels and performed the above-mentioned classification procedure, including the feature selection and classification, and a null distribution of classification accuracies was obtained. The p-value was estimated as the proportion of the accuracies in the null distribution that were greater than or equal to the actual classification accuracy.

For NSSI and NSSI+SA groups, we extracted the mean intra- and inter-modular discriminating FC strength, and the partial Pearson’s correlation analyses (age and gender were included as the confounding factors) were performed between these FC measures and clinical parameters.

## RESULTS

### Demographic and clinical characteristics

There were no significant group differences in age, gender, and education. No significant difference was observed for self-reported severity of depression and anxiety symptoms. Regarding the descriptive characteristics of NSSI, the NSSI+SA scored higher on the NSSI frequency and attention seeking of NSSI function.

**Table 1.** Demographic and clinical characteristics of the participants.

| Item | Control | NSSI | NSSI+SA | P-value |
| --- | --- | --- | --- | --- |
|  | N = 34 | N = 39 | N = 36 |  |
| Gender (M/F) | 9/25 | 6/33 | 5/31 | 0.332<sup>a</sup> |
| Age (years) | 15.65±1.61 | 15.38±1.63 | 15.06±2.08 | 0.385<sup>b</sup> |
| Edu (years) | 10.06±1.70 | 9.82±1.78 | 9.44±2.42 | 0.430<sup>b</sup> |
| PHQ scores | 15.53±6.33 | 17.64±5.88 | 18.94±5.51 | 0.056<sup>b</sup> |
| GAD scores | 11.41±5.76 | 12.79±5.26 | 13.06±5.72 | 0.419<sup>b</sup> |
| NSSI frequency (last year) | — | 12.00±9.41 | 8.00±6.02 | 0.034<sup>c*</sup> |
| NSSI function (emotion regulation) | — | 1.62±0.78 | 1.82±0.87 | 0.295<sup>c</sup> |
| NSSI function (attention seeking) | — | 0.41±0.55 | 0.75±0.67 | 0.018<sup>c*</sup> |
| NSSI function (social avoidance) | — | 0.80±0.72 | 0.90±0.85 | 0.577<sup>c</sup> |
| Life time suicide attempts | — | — | 5.53±10.86 | — |
| ERQ-CA reappraisal | 17.68±5.19 | 15.59±4.16 | 16.39±4.96 | 0.178<sup>b</sup> |
| ERQ-CA suppression | 13.35±3.04 | 14.31±2.35 | 14.92±3.11 | 0.073<sup>b</sup> |
| Current psychopharmacotherapy | 0 | 1(2.6%) | 1(2.8%) | 0.393<sup>a</sup> |
| Head movements (FD Jenkinson) | 0.068±0.023 | 0.073±0.029 | 0.075±0.031 | 0.599<sup>b</sup> |

Values represent mean (SD) or n (%) unless otherwise indicated.
NSSI, non-suicidal self-injury; PHQ, Patient Health Questionnaire; GAD: Generalized Anxiety Disorder; ERQ-CA: Emotion Regulation Questionnaire for Children and Adolescents; a, chi-square test; b, one-way ANOVA; c, two-sample t-test.
\*p < 0.05.

### Classification results based on resting-state FC

Different groups of patients were classified using linear SVM classifiers based on resting-state FC. Feature sets of different sizes were utilized, and the optimal size yielding the highest classification performances was determined (as shown in Figure 1). Accordingly, 1520 discriminating FCs were finally extracted for the distinction between NSSI and Control, 2280 discriminating FCs were finally extracted for the distinction between NSSI+SA and Control, and 320 discriminating FCs were finally extracted for the distinction between NSSI+SA and NSSI. These results highlight the discriminative power of resting-state FC in distinguishing between the different patient groups.

**Fig. 1.**
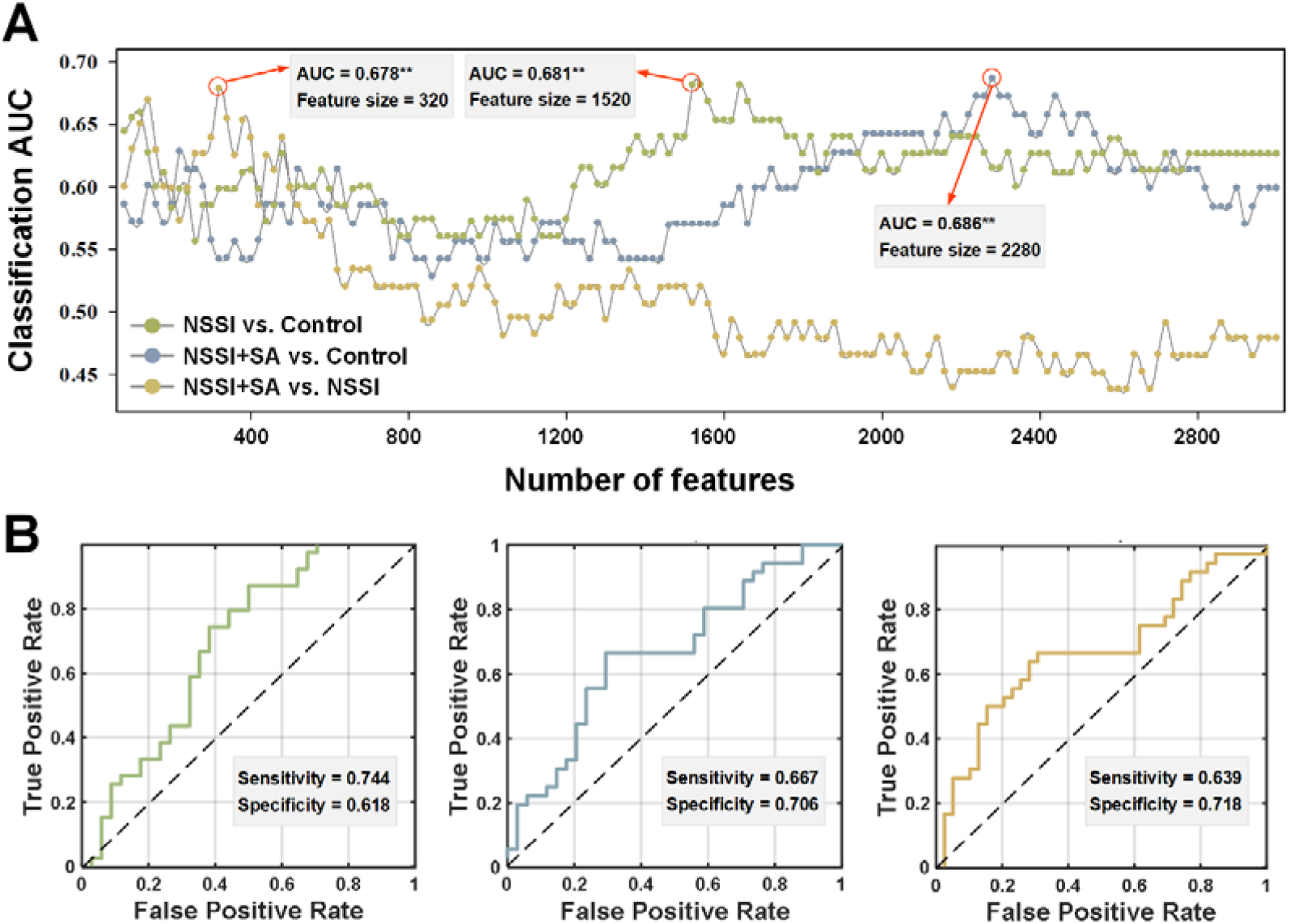
Classification performance. (A) Each curve corresponds to the classification between different groups and the maximum value was selected (**p<0.005). (B) The corresponding ROC curves.

### Group differences in resting-state FC patterns

Group differences across the entire functional connectome between groups were identified through SVM-based classification. Our focus was on the modular architecture of these discriminating features at the brain location level. Figure 2 shows the widely-distributed alternations of intra- and inter-modular resting-state FC derived from the binary classification analysis. Specifically, the matrices on the left (Figure 2A, C, and E) show the distribution of intra- and inter-modular resting-state FC with greater discriminating weights. Further, we applied the threshold of FC densities and obtained the most discriminating intra- and inter-modular resting-state FC with both greater weights and densities (the circular plots in Figure 2B, D, and F).

Compared to clinical controls, the NSSI group exhibited relatively increased resting-state FCs in the Limbic-Control, Limbic-SalVAtten, and decreased resting-state FCs in the Control, SalVAttn, Control-SalVAttn, Limbic-Default, and Limbic-DorsAttn. For adolescents engaged in both NSSI and SA, similar alternations were observed in the Limbic-Control, Limbic-SalVAtten, SalVAttn, and Limbic-Default. Different from the pure NSSI group, increased FCs were observed in Limbic-DorsAttn, Control-Visual, SalVAttn-Visual, and decreased FCs were observed in Limbic-Visual, Limbic, DorsAttn-SalVAttn. When directly comparing the NSSI+SA group to the NSSI group, discriminating FCs were predominantly observed in the Subcortical, DMN-Limbic, and Subcortical-Limbic.

**Fig. 2.**
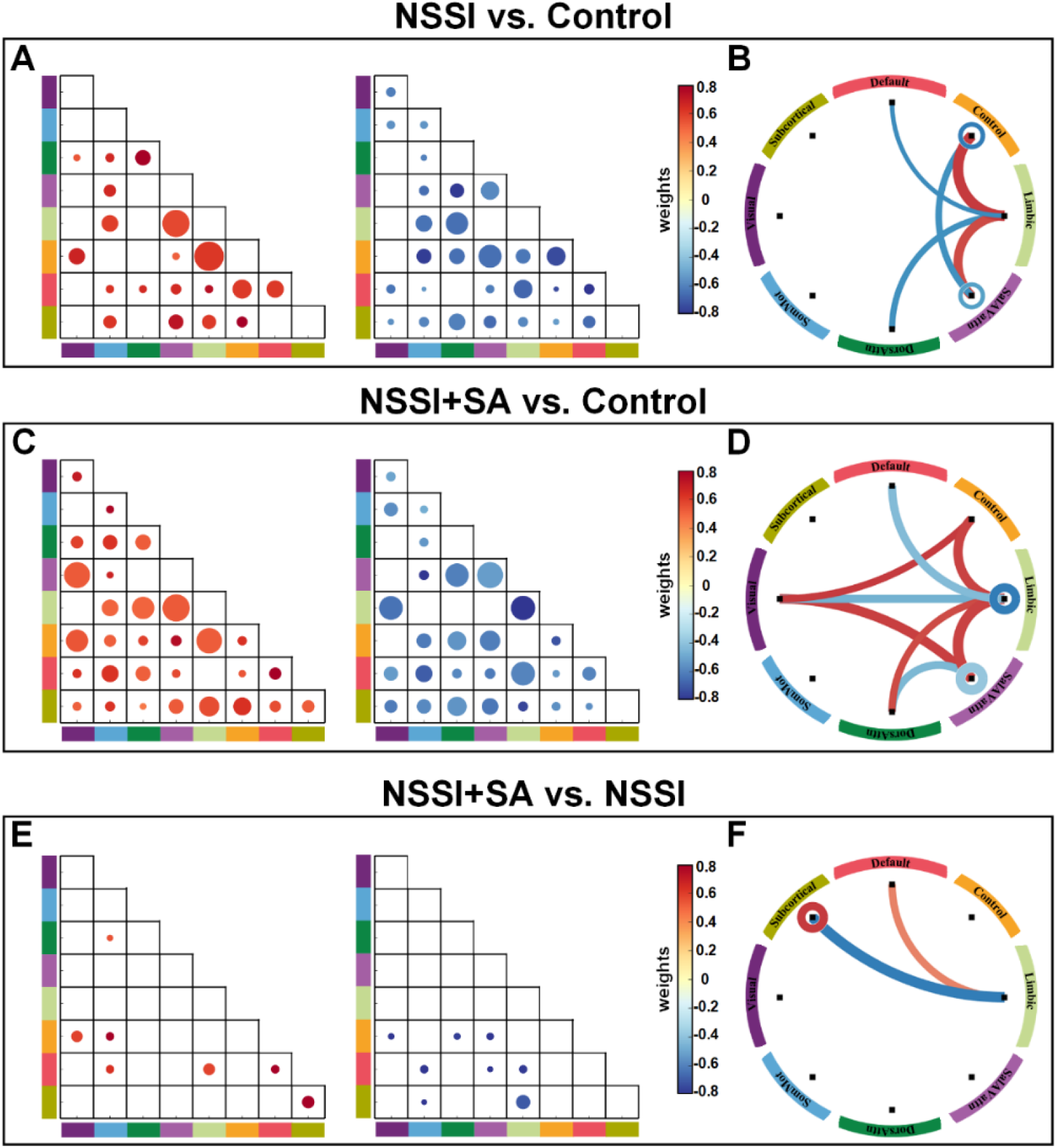
The selected resting-state functional connections contributed to the classification between every two groups. The matrices on the left (A, C, and E) show the group-mean classification weights. The size of the dots in the matrices is in proportion to the intra- and inter-modular mean discriminating weight and the scale of enlargement is the same for all matrices. The circular graphs in the middle (B and E) show the modular architecture of these consensus features with a greater percentage of connections within each module and between pairs of modules. Dots (in the matrices) and lines (in the circular graphs) with warm color represent positive discriminating weights and the reverse for dots or lines with cold color. The line thickness is in proportion to the percentage of discriminating functional connections.

The 3D glass brain representations in Figure 3 (listed in the supplementary table S1) illustrate the brain regions that contributed significantly to the classification between groups. When comparing the NSSI with the clinical Control, brain regions with positive weights located predominantly in the insular cortex (SalVAttn), orbital frontal cortex and temporal pole (Limbic), and prefrontal cortex (Control); brain regions with negative weights located predominantly in the frontal eye fields (DorsAttn), superior temporal gyrus and insular cortex (SalVentAttn), temporal pole (Limbic), cingulate and supramarginal gyrus (Control), and middle frontal cortex and precuneus (Default). When comparing the NSSI+SA with the clinical Control, brain regions with positive weights located predominantly in the frontal eye fields, superior parietal gyrus, postcentral gyrus and middle temporal gyrus (DorsAttn), insular cortex and prefrontal cortex (SalVentAttn), orbital frontal cortex and temporal pole (Limbic), cingulate and prefrontal cortex (Control), occipital cortex (Visual); brain regions with negative weights located predominantly in the frontal eye fields and inferior parietal gyrus (DorsAttn), insular cortex and parietal operculum (SalVentAttn), the temporal pole (Limbic), the medial prefrontal cortex and middle temporal gyrus (Default), the middle occipital cortex, calcarine, fusiform gyrus (Visual). When comparing the NSSI+SA with the NSSI, brain regions with positive weights located in the orbitofrontal cortex (Limbic), the posterior parietal gyrus (Default), the globus pallidus and caudate (Subcortical); brain regions associated with negative weights located in the temporal pole and orbitofrontal cortex (Limbic).

**Fig. 3.**
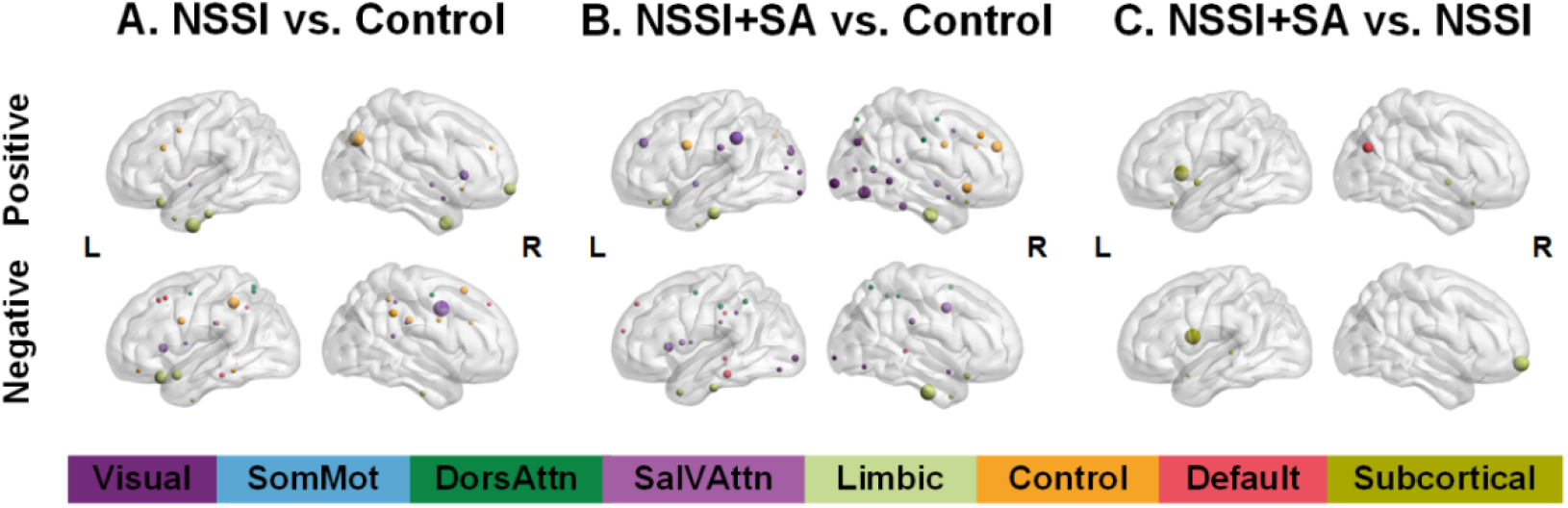
Brain regions which contributed more to the classification between the NSSI and Control (left panel), between the NSSI+SA and Control (middle panel), and between the NSSI+SA and NSSI (right panel). The colors of nodes represented the different modules of brain functional networks and the node size is in proportion to the mean discriminating weight.

### Correlation between discriminating FC strength and clinical parameters

In our analysis of the NSSI and NSSI+SA groups, we explored the correlation between altered resting-state FCs (as shown in Figure S3) and clinical variables, considering both a reference to the Control group and a direct comparison between the two patient groups. Based on the discriminating FCs extracted from the classification analysis between NSSI+SA and Control, the increased resting-state FC of Limbic-DorAttn (R = 0.434, P = 0.010) and Limbic-Visual (R = 0.362, P = 0.035) in the NSSI+SA group showed a significant positive correlation with PHQ scores (Figure 4). No significant correlation was observed between the discriminating FC and any of the clinical parameters for the NSSI group.

**Fig. 4.**
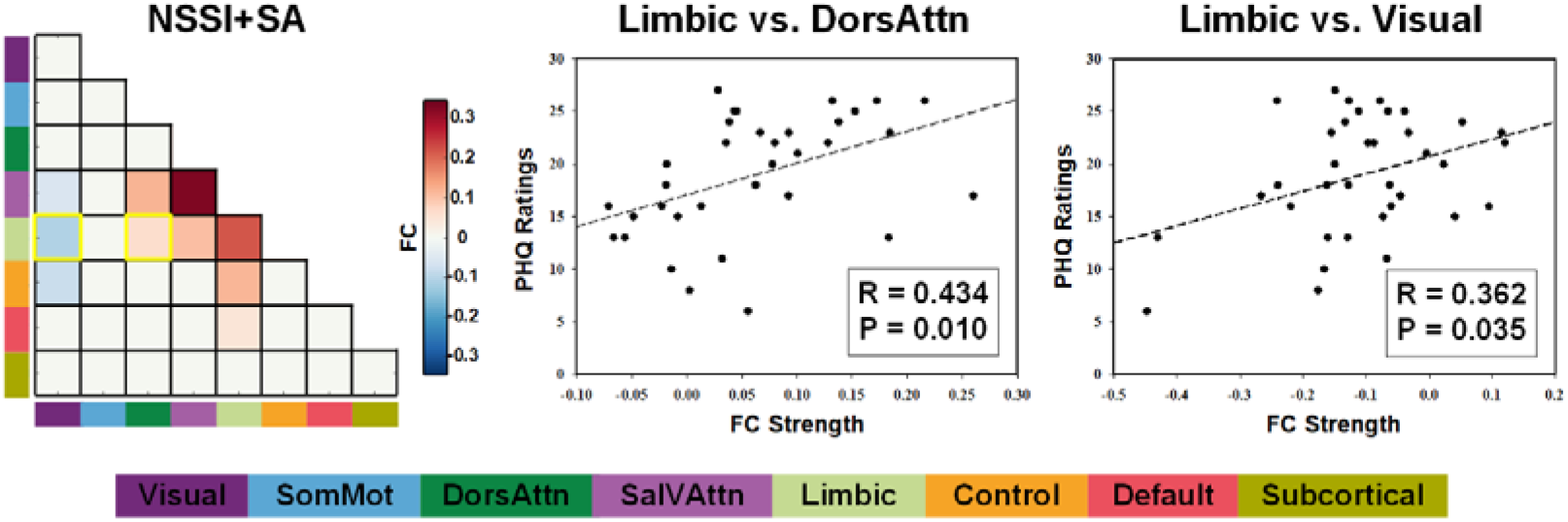
Correlation between the discriminating FC strength and clinical variables.

## DISCUSSION

Existing evidence suggests that the predisposition to suicidal and non-suicidal behaviors in adolescents is at least partly mediated by neurobiological factors. The altered cerebral function might play a key biological role in NSSI and SA of depressed adolescents [15–17, 39]. The present study was the first to examine both convergent and divergent neurobiological alternations that underpin adolescent NSSI with or without a history of SA. Compared to the clinical control group, we identified widespread aberrant intrinsic network FC patterns in the NSSI group. These alternations were not only between the three core neurocognitive intrinsic brain networks (the SalVAttn, Control, and Default networks) proposed as the triple network model of psychopathology [40], but also between these core neurocognitive intrinsic brain networks and other networks. These findings suggest that NSSI in depressed adolescents is associated with disruptions in the coordination and communication between different brain networks involved in cognitive and emotional processing. Referring to the alternations of FC in the NSSI group, the alternations of intra-/inter-network among the Control, SalVAtten, Limbic, and Default networks persist in depressed adolescents with a history of both NSSI and SA. Additionally, new FC alternations emerged, particularly including inter-network connections among the Limbic, DorsAttn, and Visual networks. These findings suggest that the combination of NSSI and SA may further influence the connectivity patterns within and between specific brain networks. Our results also expand on the link between emotional regulation deficits and suicidal behaviors by demonstrating the difference of resting-state FC between NSSI and NSSI+SA related to the Subcortical and Limbic networks. Besides, the severity of depressive symptoms only showed a significant correlation with altered FCs in Limbic-DorAttn and Limbic-Visual, strengthening the fact that these observed alternations could not all be explained by increased. We discuss potential interpretations of these findings below.

Our focus on networks is important, not only because individual brain regions do not function in isolation, but also because the network maturation is a central feature of brain development during adolescence [41, 42]. Studies of normative development have shown that functional brain networks increasingly segregate through adolescence [43], making it a vulnerable period for emotional problems [44]. Importantly, the networks that support these processes continue to mature in adolescence [41, 42] and are aberrant in depressed adolescents [45]. Our account of NSSI in adolescence is supported by a substantial body of developmental cognitive and affective neuroscience [46]. According to existing evidence using predefined ROIs, the abnormal FC tended to be spatially distributed, highlighting the importance of considering whole-brain functional networks. Menon *et al.* critically reviewed the fMRI literature and proposed a triple-network model (SalVAttn, Control, and Default network) for neuropsychiatric disorders and these three networks are associated with emotional regulation and cognitive processing [40]. NSSI or SA is a maladaptive strategy that stems from greater difficulty with emotional awareness and regulation [47]. Specific patterns of network dysfunction of core neurocognitive networks may contribute to core deficits in cognitive and affective functioning that are believed to underlie these behaviors.

The Affective Network (AN) and the Ventral Attention Network (VAN), collectively referred to as the SalVAttn network, are key brain networks involved in emotion processing and monitoring salient events [18]. The SalVAttn network, primarily centered in the anterior insular and cingulate cortex, plays a crucial role in identifying biologically and cognitively salient events that are essential for directing attention and guiding goal-directed behaviors [48]. In the NSSI compared with the Control, we found decreased intra-modular FCs of the SalVAttn and inter-network FCs between the SalVattn and Control networks, and increased FCs between the SalVattn and Limbic networks. Increased sensitivity to socioaffective pain contributes to the onset of NSSI in adolescence and it is largely believed that it is a product of an imbalance between elevated response to affective information in the environment by the SalVAttn network [49, 50], and blunted regulation by the Control network associated with effortful emotion regulation and cognitive control [51]. Besides, the Salience network also plays a critical role in processing emotion in connection with the Limbic network [48]. Therefore, increased FC between the SalVAttn and Limbic networks could be explained by the excessive emotional responses to salient external stimuli (i.e., behaviorally relevant and potentially threatening), and SalVAttn-Control hypoconnectivity may represent difficulty in the regulation of negative affect in adolescents with NSSI.

Adolescents with NSSI also showed alternations in inter-network FCs of Limbic with the Control, Default, and DorsAttn networks. The limbic network has two main nodes including the orbitofrontal cortex (Limbic-OFC) and temporal pole (Limbic-TempPole) [18]. OFC is a key region for integrating emotional information and is associated with emotional regulation in emotional disorders. In NSSI participants, greater activation was found in the OFC during the emotional processing task [52–55]. As part of an extended limbic system, the temporal pole has some role in both social and emotional processes, and dysfunction of the temporal pole has been observed in clinical disorders with socioemotional regulation [56]. The Control network is involved in the top-down regulation of attention and emotion [57], and the Default [58] and DorsAttn [59] networks are involved in internally or externally oriented attention, respectively. Therefore, our results here are in congruent with the deficits in the socioemotional regulation associated with NSSI.

The relationship between NSSI and SA is complicated and both of them lie along a spectrum of self-destructive actions [28]. Initial findings support the view that NSSI and SA in adolescents are characterized by abnormal resting-state FCs, but inconsistent patterns of alterations were reported for adolescents with NSSI and SA before [16]. NSSI is a key risk factor for future SAs, and some of the same neurobiological systems have been implicated in both NSSI and SA. According to our results, both NSSI and NSSI+SA groups showed increased FC in Limbic-Control, and Limbic-SalVAttn, and decreased FC in SalVAttn and Limbic-Default. These observations emphasize theories about a potential continuum of self-destructiveness [60] and indicates that indeed there is a potential neurobiological underpinning of the continuum from self-injury to suicidal behavior. Furthermore, it has been suggested that individuals who engage in both NSSI and SA may present with more complex psychopathology [61]. In addition to the above-mentioned convergent alternations, the divergent alternations for adolescents who engage in both NSSI and SA were observed involving the dorsal Visual network, further emphasizing the complexity of their neural connectivity patterns. In the context of NSSI, adolescents show difficulties with social cue interpretation and deficits in interpersonal problem-solving, and they are prone to negatively interpreting social situations [62]. There have been consistent findings that adolescents with repeated NSSI who additionally reported a history of SAs presented greater severity of psychopathology such as a lower level of functioning compared to those with NSSI only [12]. During the maintenance of NSSI, heightened negative bias may contribute to the development of SA [61]. Hence, our findings of alternation FC associated with the Visual network in the NSSI+SA group could be interpreted from an “information processing” perspective [63, 64]. Vision is an important part of the selective attention process, and the dorsal visual network is associated with spatial awareness and guidance of actions [65]. Hence, the processing bias in adolescents with NSSI+SA may be initiated as a perceptual visual bias, and then eventually cause a series of cognitive and affective symptoms.

Furthermore, when comparing the NSSI group and the NSSI+SA group directly, both groups reported emotion regulation as their major motivation for NSSI behaviors, and the NSSI+SA group reported a higher rating for the attention-seeking factor. The stronger need for attention-seeking for adolescents with SA was in consistent with Joiner’s “interpersonal-psychological theory of suicidal behavior”, which states belongingness as one of the main predictors of suicidal behavior [60]. Emotion regulation was the most common reason for NSSI consistently suggested in many previous studies [66]. While a basic understanding of the relationship between NSSI and emotion regulation deficits has been established, one recently published study supported the possibility of unique associations between emotion regulation deficits with more frequent NSSI engagement and more suicidal attempts [67]. According to our results, the NSSI+SA group reported a higher frequency of self-injury behaviors, which means that suicidal behavior may gradually become another coping strategy due to the more pronounced habituation to pain and fear because of repeated self-injury behaviors when emotion dysregulation persists. Besides, the differences in resting-state FC between the NSSI group and the NSSI+SA group were mainly related to the basal ganglia of the Subcortical network. The basal ganglia traditionally have been assigned to roles within the motor domain, however, the basal ganglia have connections with a broad limbic network and yet recent research has proposed its contribution to a variety of limbic and cognitive processes including emotion recognition, reward- and decision-making [68]. Therefore, the current study expands on the link between emotional regulation deficits and suicidal behaviors by demonstrating the difference of resting-state FC related to the Subcortical and Limbic networks. Since the self-reported questionnaire for emotion regulation did not yield significant differences among groups, future studies could use a more objective assessment of emotion regulation function to validate the results observed here.

### Limitations and future directions

Despite the valuable insights gained from this study, there are still some limitations. First, the cross-sectional design limits our ability to establish the causal relationship or determine the development trajectory of the observed aberrant patterns. To fully comprehend the emergence and progression of these patterns, longitudinal studies that track individuals over time are needed. Secondly, the reliance on retrospective self-report to collect data on self-harm behaviors and depression symptoms introduces the potential reporting or recall biases, which may affect the accuracy and reliability of the obtained information. Future research could consider incorporating more objective measures to mitigate these biases. Thirdly, the sample of depressed adolescents limits the generalizability of the identified FC patterns to a broader population of adolescents engaging in NSSI and SA. As a related point, this study included both adolescent females and males (although mostly females). Given consistently observed sex differences in the prevalence of self-injurious behaviors [69] and brain function during adolescence [70], there may be sex differences in the neural mechanisms of NSSI and SA. To gain a more comprehensive understanding, future studies should investigate these neural circuits regarding different psychiatric conditions and potential sex differences. By addressing these limitations, we can further enhance our understanding of the neural mechanisms underlying NSSI and SA.

## CONCLUSIONS

In summary, this study represents the first attempt to explore the convergent and divergent neurobiological alternations associated with NSSI and SA in adolescents. By focusing on network-level analysis, this study extends our current understanding of the neural mechanisms underlying these behaviors at a circuit level. The findings suggest the presence of a continuum of self-destructiveness in adolescents and provide insights into the neural mechanism that contributes to this continuum. Additionally, the results can potentially aid in the identification of those at risk for SA among those engaging in NSSI and contribute to the development of neurobiologically informed interventions.

## ACKNOWLEDGEMENTS

We would like to extend our thanks to the adolescents who participated in this study and their parents. This work was supported by the National Natural Science Foundation of China (No. 62276169 and 82272114), Natural Science Foundation of Guangdong Province, China (No. 2021A1515011152 and 2023A1515010840), Shenzhen Soft Science Research Program Project (No. RKX20220705152815035), Shenzhen Science and Technology Research and Development Fund for Sustainable Development Project (No. KCXFZ20201221173400001 and KCXFZ20201221173613036), Shenzhen’s Sanming Project of Medicine (No. SZSM202111009 and SZSM202011014).

## AUTHOR CONTRIBUTIONS

Conception and design of this study: YZ, ZZ. Acquisition of data: GL, HX, XY, XL, XZ. Analysis of data: LL, ZL. Drafting of the article: LL. Critical revision of the article: ZL, YZ, ZZ. All authors approved the final version of the manuscript.

## ETHICS APPROVAL

The study protocol was approved by the Human Research Ethics Committee of Shenzhen Kangning Hospital. Informed, written consent was obtained from all participants and their caregivers prior to participation in the study.

## Supplementary materials

### 1. MATERIALS AND METHODS

#### 1.1 Participants

The Chinese version of the Functional Assessment of Self-Mutilation (FASM) [1] was used to assess methods, frequency, and functions of NSSI behaviors in the past year. Participants were asked if they had engaged in any of the 10 listed self-harm behaviors in the past 12 months. Participants who answered “yes” were NSSI+ while those who answered “yes” were NSSI-. The reported frequency of NSSI behaviors in the last year was converted to ratings on a 5-point scale (0: never; 1: 0-25%; 2: 25%-50%; 3: 50%-75%; 4: >75%). Besides, participants were asked if they had tried to take their life before. Participants who answered “no” were SA-while participants who answered “yes” were SA+ and made a further report about times of suicidal behavior. The Chinese version of the FASM uses a three-factor model, named emotion regulation, attention seeking, and social avoidance, which gives rise to a reliable structure of the NSSI function in the Chinese cultural context. Scores for items belonging to each factor were summed to obtain the scores for each factor. To make the endorsement of the three factors comparable, the scores were scaled by dividing the summed scores by the number of subscales for each factor.

#### 1.2 Data acquisition and preprocessing

MRI scanning was performed at the Department of Radiology at Shenzhen Kangning Hospital using a 3.0-Tesla scanner MRI (Prisma, Siemens, Germany). Resting-state fMRI was collected using an echo-planar imaging (EPI) sequence with the following parameters: TR/TE = 2000/30 ms; flip angle = 90L; slice thickness = 2.5 mm; acquisition matrix = 88×88; number of slices = 58; and 240 time points. During the scanning, each participant was asked to keep still with their eyes closed, and not to think about anything or fall asleep. High-resolution three-dimensional brain volume T1-weighted imaging was collected using the following parameters: TR/TE/IT = 2530/2.27/1100 ms; flip angle = 7L; slice thickness = 1 mm; acquisition matrix = 256×256; number of slices = 144. The preprocessing of resting-state fMRI was performed using Data Processing Assistant for Resting-State fMRI Advanced Edition V2.2 (DPARSFA, http://rfmri.org/DPARSF) toolkit with Statistical Parametric Mapping (SPM8, https://www.fil.ion.ucl.ac.uk/spm/) in MATLAB (R2010b) (MathWorks, Natick, MA, USA).

The main preprocessing steps were as follows: (1) The first 5 volumes were removed to allow for signal equilibration; (2) slice-timing correction was performed and the middle slice was used as the reference slice; (3) motion correction realignment was performed; five participants with excessive head motion, as operationalized by having a mean FD_Jenkinson_>0.2mm, were excluded; (4) Spatial normalization was performed to the Montreal National Institute space (MNI) with a voxel size of 3×3×3mm^3^; (5) to reduce the effect of physiological noise, a series of nuisance covariates were regressed out using the Friston-24 model [2], including the mean signal of the white matter, cerebrospinal fluid signal, and head motion parameters; (6) temporal bandpass filtering (0.01-0.1 Hz) was further performed to reduce low-frequency drift and high-frequency noise; (7) spatial smoothing was performing using a 4 mm full-width at high maximum (FWHM) Gaussian kernel. Additionally, to ensure that residual motion was not a confound, mean framewise displacement was compared between groups, and no significant effect of motion was observed (one-way ANOVA, P = 0.599).

#### 1.3 Functional network construction

To define network nodes, human brains were parcellated into 232 cortical and subcortical regions of interest (ROIs). The 200 Cortical ROIs were obtained from the scale-200 version of the whole-brain functional parcellation of Schaefer et al [3]. The 32 subcortical ROIs were obtained from a recent subcortical functional parcellation [4]. The 232 ROIs can be assigned into eight intrinsic connectivity networks (ICNs): (1) visual network, (2) somatomotor network (SomMot), (3) dorsal attention network (DorsAttn), (4) salience/ventral attention network (SalVAttn), (5) limbic network, (6) control network, (7) default mode network, and (8) subcortical network [5]. We extracted the averaged BOLD time series for each of the 232 ROIs for each subject. A 232×232 correlation matrix for each subject was subsequently constructed by calculating Pearson’s correlation coefficients between the BOLD time series from all possible pairs of ROIs. A Fisher’s r-to-z transformation was further applied to the correlation matrices. The upper triangular part of the resting-state FC matrix was extracted to construct one feature vector for each subject.

**Figure S1.**
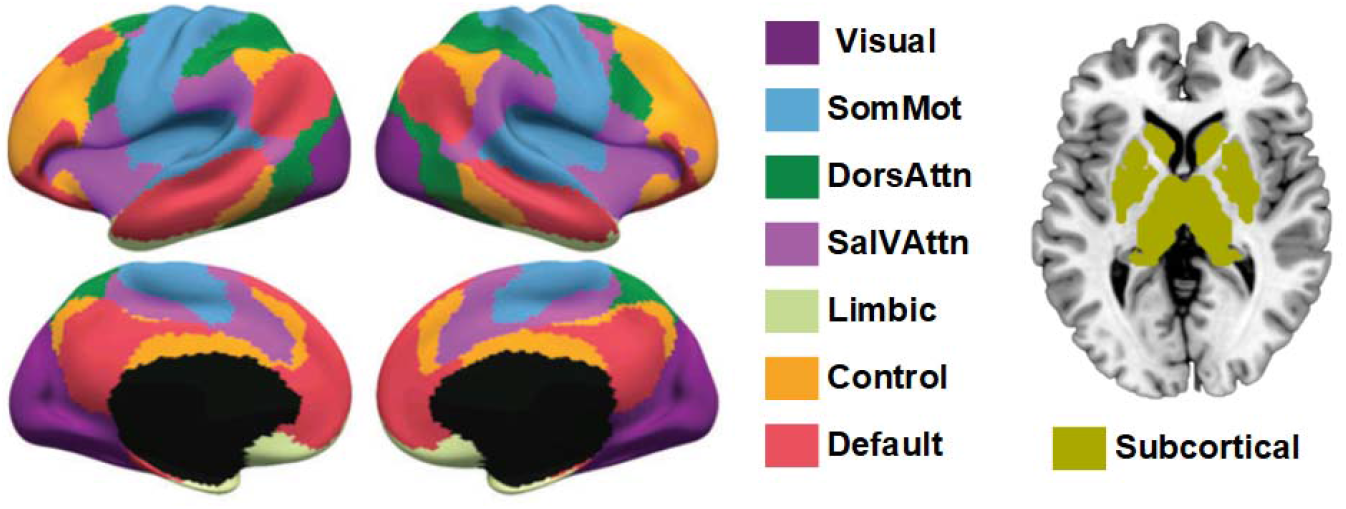
Whole-brain functional parcellation.

#### 1.4 Support vector machine training and classification

Because the feature size was much larger than the sample size, a feature selection algorithm was applied to identify discriminating features between two groups and remove unnecessary information. In this study, the F-scores method was employed for feature ranking [6]. F score is simple, generally quite effective, and used in previous resting-state fMRI studies [7]. The F score of each feature is calculated by the ratio between the variance between groups and the variances within each of the two groups. For features of topological nodal metrics, the F score was first calculated for each metric and then averaged. The larger the F-score is, the more likely the feature is to be more discriminative between the two groups.

To identify the optimally differentiating features between the two groups, nested-leave-one-out-cross-validation (LOOCV) was applied. Prior to analysis, each feature was normalized across subjects in the training sample by subtracting the mean value and dividing by the standard deviation, and the same mean and standard deviation were used to scale the test data. The feature number was tested from 20 to 3000 with a step of 20. For each LOOCV iteration, the data were divided into test and training samples. The F-scores of all features were calculated and then ranked within the training set. Specificity, sensitivity, and area under curve (AUC) were calculated to measure the performance of the selected classifier. The prediction performance was calculated for all the LOOCV iterations of each step of feature size, and the smallest step which achieved the highest prediction AUC value was chosen.

### 2. RESULTS

#### 2.1 Classification results based on resting-state FC

Resting-state fMRI FCs that provide relevant information to distinguish between each pair of groups. Matrices in Figure S2 show the consensus features which were normalized to [-1 1] across all connections.

**Figure S2.**
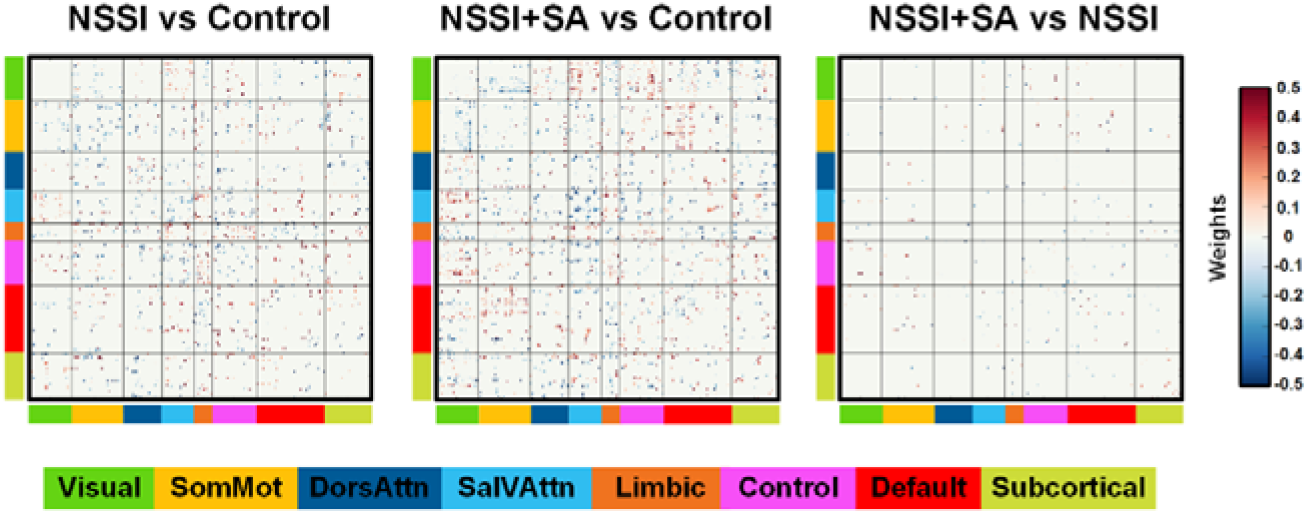
The mean weights of all consensus features.

In order to extract the most discriminating features at the network level, we first thresholded the weights of consensus features and then threshold the densities of intra- and inter-modular features. Figure S3 shows the averaged values of discriminating intra- and inter-modular features for each group.

**Figure S3.**
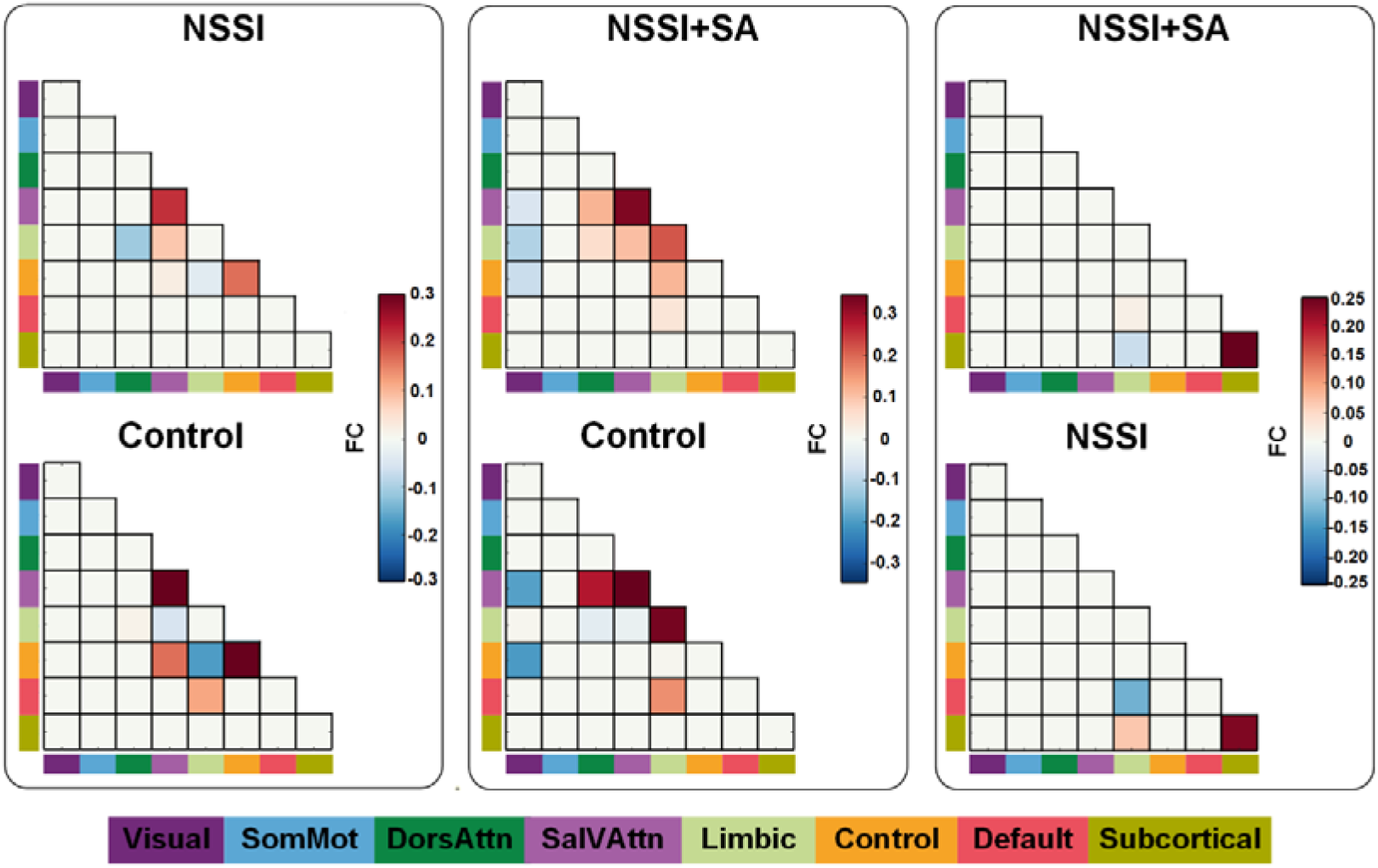
The mean resting-state FC strength changes between groups for intra- and inter-modular discriminating features.

The predictive contributions of the consensus intra- and inter-modular features with both greater classification weights and densities were also visualized at the nodal level. The brain regions which contributed more to the classification between groups are listed in the supplementary table S1. The anatomical labels and MNI coordinates were provided.

**Table S1.**
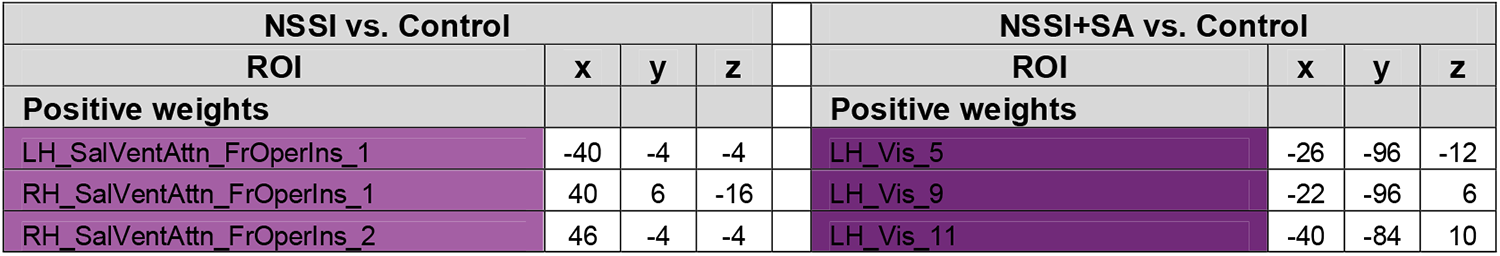
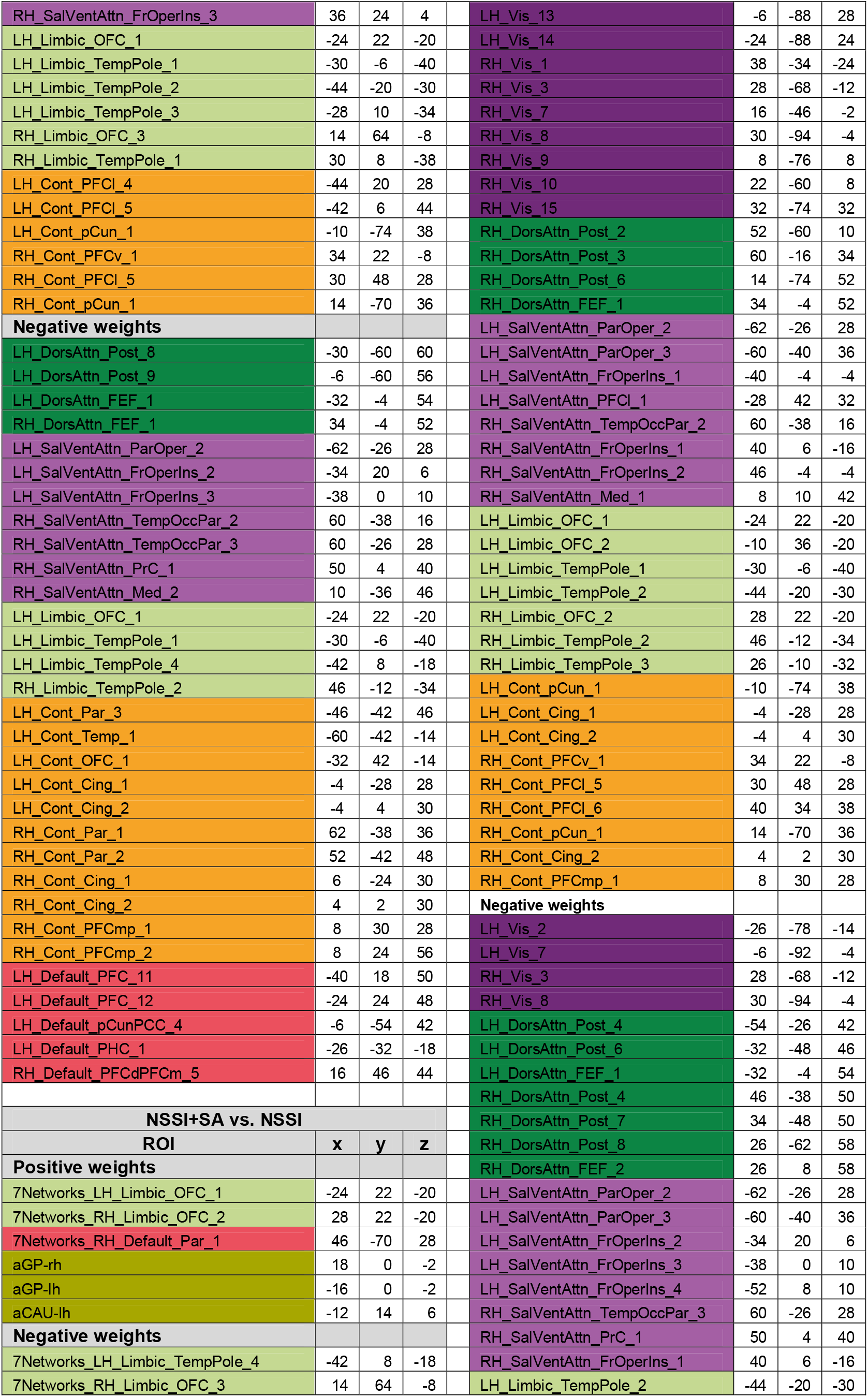
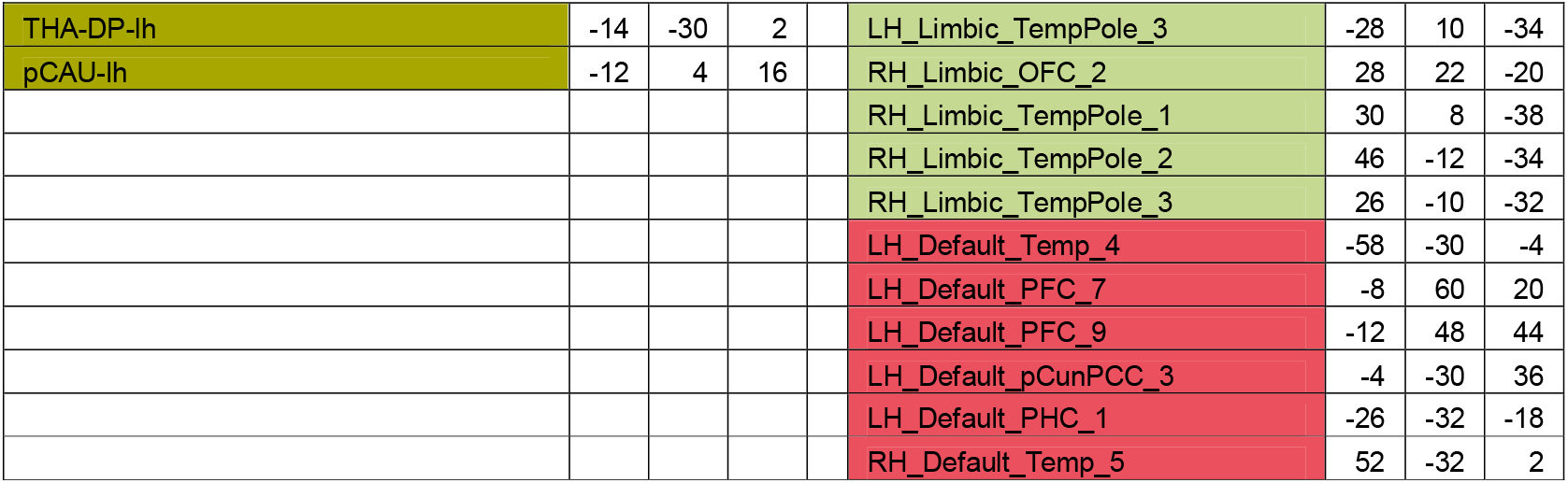
Information on brain regions contributed more to the classification between groups.

